# microbialPhenotypes: An R package that analyzes high-throughput microbial phenotype data

**DOI:** 10.1101/2020.06.29.177659

**Authors:** Peter I-Fan Wu, James C. Hu, Deborah A. Siegele

## Abstract

Various microbial high-throughput phenotyping techniques have been vastly conducted to infer functions of genes, generating large numbers of valuable datasets whose potential in providing insights to characterize genes hasn’t been fully exploited. Therefore, computational tools that allow unbiased, systematic analysis of these data also have become vital. Here we describe a package that evaluates high-throughput microbial phenotype data by one or several sets of associated functional annotations are provided. In addition, some helper functions are provided to help clean high-throughput microbial phenotype data.

## Introduction

Phenotypes play important roles in characterizing the functions of genes, particularly in microbiology, where the largest number of tests could be done much easier (Tohsato & Mori, 2008). With rising technologies (Kritikos et al., 2017; Nichols et al., 2011; Wetmore et al., 2015), high-throughput experimental approaches that measure large number of phenotypes under various conditions have flourished with complementary statistical methods in querying the behavior of gene products (Collins, Schuldiner, Krogan, & Weissman, 2006; Nichols et al., 2011; Price et al., 2018; Rishi et al., 2020). Although there are already many computational approaches written as R packages to analyze phenotype data (Deng, Gao, Wang, & Guo, 2015; Vaas et al., 2013; Vuckovic, Gasparini, Soranzo, & Iotchkova, 2015) (Vehkala, Shubin, Connor, Thomson, & Corander, 2015), software that quickly tests the potential of such high-throughput data in interpreting gene functions is lacking. Here we have wrapped the analytical pipeline of evaluating the phenotype data using annotation sets into a package named microbialPhenotypes. In this package, we provide 8 functions that are specific in dealing with high-throughput microbial phenotype data, as well as 10 functions that are meant to be more supplementary. A high-throughput *E. coli* phenotype dataset (Nichols et al., 2011) is used for the examples provided below. We note that there is room for significant improvement when the analytical pipeline is improved. Despite that the functions provided here started from a perspective of gaining biological insights for microbial phenotype data, the potential of using them as a tool for more general purposes should not be limited. If there are data from other research domains that has a similar structure to the example described here, the utility of this package can be much more extended. For example, data from animal cells (Alonezi et al., 2016, 2017). In addition, knowing that there are resources for complicated machine learning algorithms to do classification of functions, our work here does not aim to improve those methods. Rather, it tries to quickly assess the usefulness of the newly generated high-throughput phenotype data before implementing more advanced classification schema

### Installation and functions

The **microPhenotypes** package can be downloaded from GitHub: https://github.com/peterwu19881230/microbialPhenotypes. To load the functions, source the .R files under R/.

### Input data

This package assumes the input phenotypic profiles be in flat files (e.g., csv or tsv files) in a matrix format where each row represents a mutant strain and each column is a condition where the phenotypes of the mutant strains are assayed. The values in the columns represent the phenotypes.

Users can read their data into R using either the read.csv() or read.table() function with the appropriate arguments based on the format of the file.

For example:

> phenotype_data <- read.csv(file=“my_phenotype_profile.csv”, header = TRUE)

> head(phenotype_data)

|  | STREPTOMYCIN.0.05 | SUCCINATE | SULFAMONOMETHOXINE.100 | TAUROCHOLATE.0.1 | TAUROCHOLATE.0.5 |
| --- | --- | --- | --- | --- | --- |
| ECK3997-purD | -1.48 | -12.30 | 3.33 | -13.46 | -13.79 |
| ECK0516-purE | 0.43 | -6.75 | 2.63 | -8.27 | -9.81 |
| ECK3763-ilvD | -1.41 | -11.72 | 0.34 | -0.69 | 0.32 |
| ECK3766-ilvC | -0.58 | -8.53 | -0.81 | 0.06 | 0.51 |
| ECK3762-ilvE | -0.01 | -11.93 | 0.02 | -0.68 | -0.46 |
| ECK3764-ilvA | -0.15 | -11.55 | -0.84 | -0.04 | -0.19 |

### Discretize the input data

Quantitative data can be transformed into categorical data using either the BinaryConvert() or TernaryConvert() function, which produce output in a binary (0,1) or ternary (−1,0,1) form, respectively. The threshold argument is used to specify the phenotypic score cutoff used to select strains with a phenotype that is significantly different from that of the designated control, which is usually the phenotype of the wildtype.

For example:

> ter_phenotype <- ternary_convert(matrix=phenotype_data, threshold =0.5)

> head(ter_phenotype)

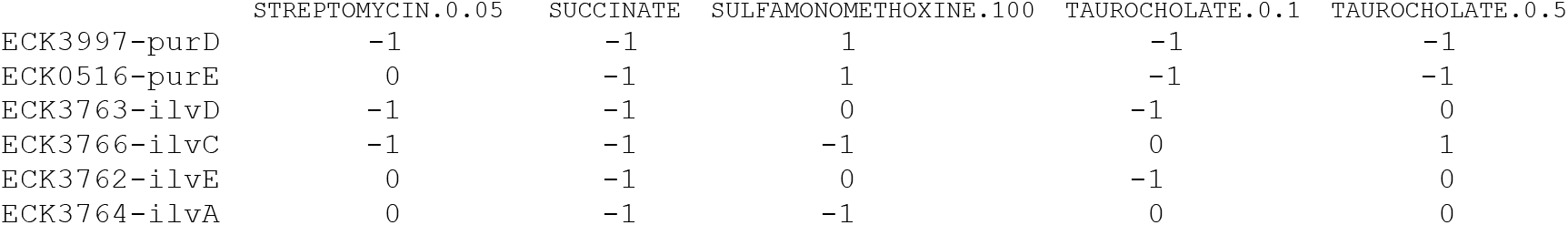

|  | STREPTOMYCIN.0.05 | SUCCINATE | SULFAMONOMETHOXINE.100 | TAUROCHOLATE.0.1 | TAUROCHOLATE.0.5 |
| --- | --- | --- | --- | --- | --- |
| ECK3997-purD | -1 | -1 | 1 | -1 | -1 |
| ECK0516-purE | 0 | -1 | 1 | -1 | -1 |
| ECK3763-ilvD | -1 | -1 | 0 | -1 | 0 |
| ECK3766-ilvC | -1 | -1 | -1 | 0 | 1 |
| ECK3762-ilvE | 0 | -1 | 0 | -1 | 0 |
| ECK3764-ilvA | 0 | -1 | -1 | 0 | 0 |

> binary_phenotype <- binary_convert (matrix=phenotype_data, threshold =0.5)

> head(binary_phenotype)

|  | STREPTOMYCIN.0.05 | SUCCINATE | SULFAMONOMETHOXINE.100 | TAUROCHOLATE.0.1 | TAUROCHOLATE.0.5 |
| --- | --- | --- | --- | --- | --- |
| ECK3997-purD | 1 | 1 | 1 | 1 | 1 |
| ECK0516-purE | 0 | 1 | 1 | 1 | 1 |
| ECK3763-ilvD | 1 | 1 | 0 | 1 | 0 |
| ECK3766-ilvC | 1 | 1 | 1 | 0 | 1 |
| ECK3762-ilvE | 0 | 1 | 0 | 1 | 0 |
| ECK3764-ilvA | 0 | 1 | 1 | 0 | 0 |

### Calculate similarities/distances between phenotypic profiles

The pairwise similarity/distance between phenotypic profiles can be calculated by a variety of functions, such as Hamming distance, Pearson Correlation Coefficient, Spearman Correlation Coefficient, Mutual Information. The method selected will depend on the type of assay used.

The function implemented in this package is hamming distance:

> hamming_dist=hamming_distance(head(ter_phenotype))

> hamming_dist

|  | ECK3997-purD | ECK0516-purE | ECK3763-ilvD | ECK3766-ilvC | ECK3762-ilvE | ECK3764-ilvA |
| --- | --- | --- | --- | --- | --- | --- |
| ECK3997-purD | 0 | 1 | 2 | 3 | 3 | 4 |
| ECK0516-purE | 1 | 0 | 3 | 4 | 2 | 3 |
| ECK3763-ilvD | 2 | 3 | 0 | 3 | 1 | 3 |
| ECK3766-ilvC | 3 | 4 | 3 | 0 | 4 | 2 |
| ECK3762-ilvE | 3 | 2 | 1 | 4 | 0 | 2 |
| ECK3764-ilvA | 4 | 3 | 3 | 2 | 2 | 0 |

Example for a self-defined function not included in this package. Pearson Correlation Coefficient was chosen because it’s widely used in high-throughput microbial phenotype data:

> pearson_dist=1- cor(t(phenotype_data),method=“pearson”)

> pearson_dist

|  | ECK3997-purD | ECK0516-purE | ECK3763-ilvD | ECK3766-ilvC | ECK3762-ilvE | ECK3764-ilvA |
| --- | --- | --- | --- | --- | --- | --- |
| ECK3997-purD | 0.00000000 | 0.01130833 | 0.67443135 | 0.792573199 | 0.614310165 | 0.709983271 |
| ECK0516-purE | 0.01130833 | 0.00000000 | 0.78293109 | 0.891990335 | 0.706927257 | 0.800223580 |
| ECK3763-ilvD | 0.67443135 | 0.78293109 | 0.00000000 | 0.015107990 | 0.012529945 | 0.017520039 |
| ECK3766-ilvC | 0.79257320 | 0.89199034 | 0.01510799 | 0.000000000 | 0.017882605 | 0.006075217 |
| ECK3762-ilvE | 0.61431017 | 0.70692726 | 0.01252994 | 0.017882605 | 0.000000000 | 0.005903022 |
| ECK3764-ilvA | 0.70998327 | 0.80022358 | 0.01752004 | 0.006075217 | 0.005903022 | 0.000000000 |

### Parse gene annotations

In order to link the functional annotations associated with a gene to the phenotypic profile of a strain where that gene is mutated, three functions, attr_list(), one_attr() and generate_pairs_similarity_coannotation (), are provided where:

attr_list() takes in a table that has mapping from mutant ids to their annotations and form a list from such mapping, for example, if starting from inputting the strain names and corresponding annotations:

attr_list() creates a list takes where each strain is associated with all the functional annotations and creates a list. To illustrate this, we will create a table with the name “name_attribute”:

> name_attribute = data.frame(id=c(“ECK3762-ilvE”, “ECK3762-ilvE”, “ECK3762-ilvE”, “ECK3762-ilvE”, “ECK3762-ilvE”, “ECK3762-ilvE”, “ECK3762-ilvE”, “ECK3762-ilvE”, “ECK3762-ilvE”, “ECK3762-ilvE”, “ECK3764-ilvA”, “ECK3764-ilvA”, “ECK3764-ilvA”, “ECK3997-purD”, “ECK3997-purD”, “ECK3997-purD”, “ECK3997-purD”, “ECK3997-purD”, “ECK0516-purE”, “ECK0516-purE”, “ECK0516-purE”, “ECK3763-ilvD”, “ECK3763-ilvD”, “ECK3763-ilvD”, “ECK3763-ilvD”, “ECK3766-ilvC”, “ECK3766-ilvC”, “ECK3766-ilvC”, “ECK3766-ilvC”, “ECK3766-ilvC”, “ECK3766-ilvC”), annot=c(“ALANINE-VALINESYN-PWY”, “THREOCAT-PWY”, “ALL-CHORISMATE-PWY”, “PWY0-1061”, “PHESYN”, “ILEUSYN-PWY”, “LEUSYN-PWY”, “VALSYN-PWY”, “COMPLETE-ARO-PWY”, “BRANCHED-CHAIN-AA-SYN-PWY”, “THREOCAT-PWY”, “ILEUSYN-PWY”, “BRANCHED-CHAIN-AA-SYN-PWY”, “PRPP-PWY”, “DENOVOPURINE2-PWY”, “PWY-6121”, “PWY-6122”, “PWY-6277”,”PRPP-PWY”, “PWY-6123”, “DENOVOPURINE2-PWY”, “THREOCAT-PWY”, “ILEUSYN-PWY”, “VALSYN-PWY”, “BRANCHED-CHAIN-AA-SYN-PWY”, “THREOCAT-PWY”, “PANTO-PWY”, “ILEUSYN-PWY”, “VALSYN-PWY”, “PANTOSYN-PWY”, “BRANCHED-CHAIN-AA-SYN-PWY”))

> name_attribute

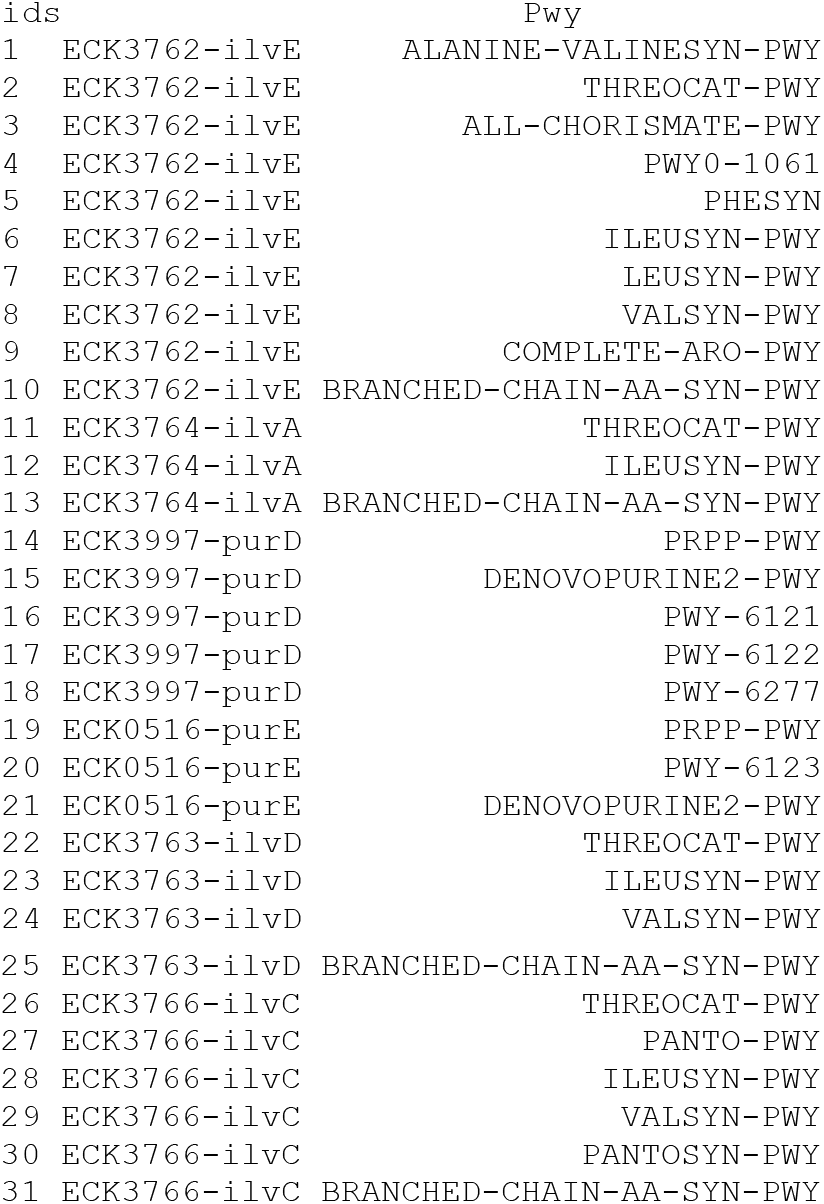

> attr_list(name_attribute)

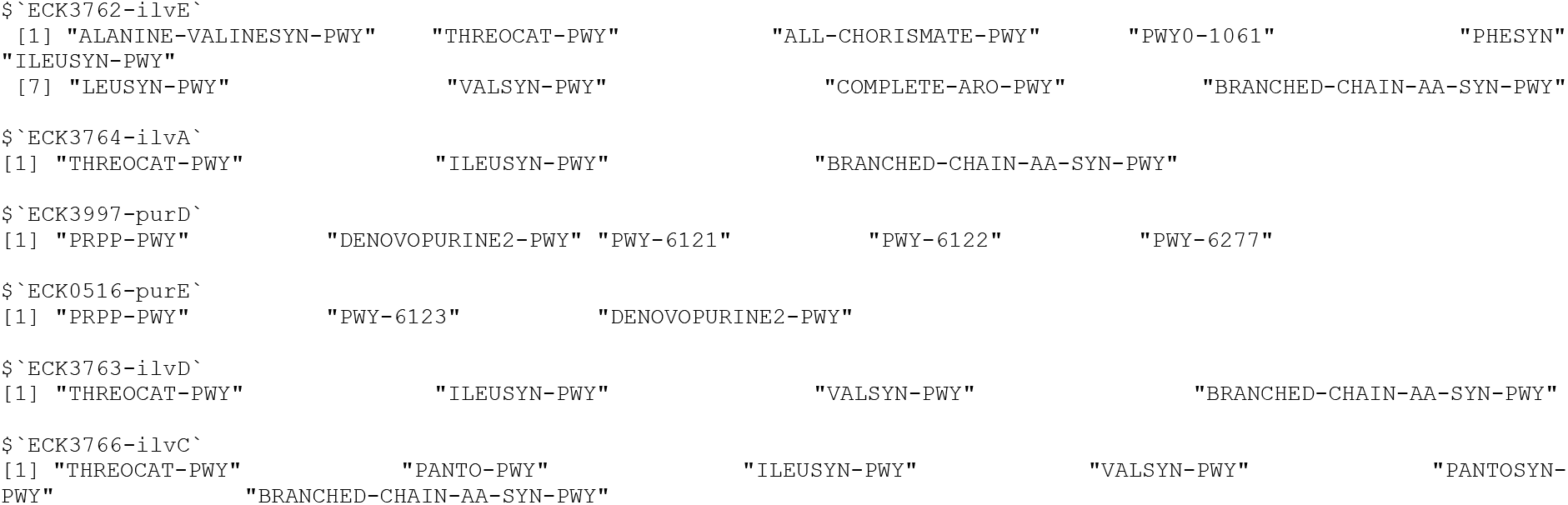

one_attr() takes the output from attr_list() as the input. It generates a table that contains all possible combination of mutants and whether they share annotations (0 stands for not having any same annotations, 1 for having at least 1 same annotation). For example:

> attribute_list=attr_list(name_attribute)

> one_attr(attribute_list)

|  | mutant1 | mutant2 | sameORnot |
| --- | --- | --- | --- |
| 1 | ECK3762-ilvE | ECK3764-ilvA | 1 |
| 2 | ECK3762-ilvE | ECK3997-purD | 0 |
| 3 | ECK3762-ilvE | ECK0516-purE | 0 |
| 4 | ECK3762-ilvE | ECK3763-ilvD | 1 |
| 5 | ECK3762-ilvE | ECK3766-ilvC | 1 |
| 6 | ECK3764-ilvA | ECK3997-purD | 0 |
| 7 | ECK3764-ilvA | ECK0516-purE | 0 |
| 8 | ECK3764-ilvA | ECK3763-ilvD | 1 |
| 9 | ECK3764-ilvA | ECK3766-ilvC | 1 |
| 10 | ECK3997-purD | ECK0516-purE | 1 |
| 11 | ECK3997-purD | ECK3763-ilvD | 0 |
| 12 | ECK3997-purD | ECK3766-ilvC | 0 |
| 13 | ECK0516-purE | ECK3763-ilvD | 0 |
| 14 | ECK0516-purE | ECK3766-ilvC | 0 |
| 15 | ECK3763-ilvD | ECK3766-ilvC | 1 |

generate_pairs_similarity_coannotation() generates the pairwise similarity table that contains the strain pairs, similarity/distance value and a Boolean column of whether the corresponding strain pairs share the same annotation(s). It takes phenotype data, the result from attr_list() and a function as an argument to specify the similarity metric. For example:

> attribute_list<-one_attr(name_attribute)

> names(attribute_list)<-rownames(phenotype_data) #(Must do) Synchronize the strain names

> generate_pairs_similarity_coannotation(data= phenotype_data,attribute_list=attribute_list,

+ dist_metric=pcc_dist)

| mutant1 | mutant2 | similarity | same_annot |  |
| --- | --- | --- | --- | --- |
| 1 | ECK3762-ilvE | ECK3764-ilvA | 0.005903022 | 1 |
| 2 | ECK3766-ilvC | ECK3764-ilvA | 0.006075217 | 1 |
| 3 | ECK3997-purD | ECK0516-purE | 0.011308326 | 1 |
| 4 | ECK3763-ilvD | ECK3762-ilvE | 0.012529945 | 1 |
| 5 | ECK3763-ilvD | ECK3766-ilvC | 0.015107990 | 1 |
| 6 | ECK3763-ilvD | ECK3764-ilvA | 0.017520039 | 1 |
| 7 | ECK3766-ilvC | ECK3762-ilvE | 0.017882605 | 1 |
| 8 | ECK3997-purD | ECK3762-ilvE | 0.614310165 | 0 |
| 9 | ECK3997-purD | ECK3763-ilvD | 0.674431348 | 0 |
| 10 | ECK0516-purE | ECK3762-ilvE | 0.706927257 | 0 |
| 11 | ECK3997-purD | ECK3764-ilvA | 0.709983271 | 0 |
| 12 | ECK0516-purE | ECK3763-ilvD | 0.782931085 | 0 |
| 13 | ECK3997-purD | ECK3766-ilvC | 0.792573199 | 0 |
| 14 | ECK0516-purE | ECK3764-ilvA | 0.800223580 | 0 |
| 15 | ECK0516-purE | ECK3766-ilvC | 0.891990335 | 0 |

, where pcc_dist = 1-cor(t(your_data),method=“pearson”)

### Get metrics derived of the confusion matrix

After the similarity and co-annotation columns are computed as above, get_confusionMatrix() can be used to get the similarity-based confusion matrices and the derived metrics: sensitivity, specificity, precision, and accuracy. get_confusionMatrix() also permutate the co-annotation column and calculate the confusion matrices and other derived metrics as negative controls. For example:

> new <-generate_pairs_similarity_coannotation(data= phenotype_data, attribute_list=attribute_list, dist_metric=pcc_dist)

> get_confusionMatrix_and_metrics (df=new, annot=“same_annot”,similarity=“similarity”)

| TP | FP | TN | FN | sensitivity | specificity | precision | accuracy | random_TP | random_FP | random_TN | random_FN |
| --- | --- | --- | --- | --- | --- | --- | --- | --- | --- | --- | --- |
| random_sensitivity | random_specificity | random_precision | random_accuracy |  |  |  |  |  |  |  |  |
| 1 | 1 | 0 | 8 | 6 | 0.1428571 | 1.000 | 1.0000000 | 0.6000000 | 0 | 1 | 7 |
| 7 |  |  |  |  | 0.0000000 | 0.875 | 0.0000000 | 0.4666667 |  |  |  |
| 2 | 2 | 0 | 8 | 5 | 0.2857143 | 1.000 | 1.0000000 | 0.6666667 | 0 | 2 | 6 |
| 7 |  |  |  |  | 0.0000000 | 0.750 | 0.0000000 | 0.4000000 |  |  |  |
| 3 | 3 | 0 | 8 | 4 | 0.4285714 | 1.000 | 1.0000000 | 0.7333333 | 0 | 3 | 5 |
| 7 |  |  |  |  | 0.0000000 | 0.625 | 0.0000000 | 0.3333333 |  |  |  |
| 4 | 4 | 0 | 8 | 3 | 0.5714286 | 1.000 | 1.0000000 | 0.8000000 | 0 | 4 | 4 |
| 7 |  |  |  |  | 0.0000000 | 0.500 | 0.0000000 | 0.2666667 |  |  |  |
| 5 | 5 | 0 | 8 | 2 | 0.7142857 | 1.000 | 1.0000000 | 0.8666667 | 0 | 5 | 3 |
| 7 |  |  |  |  | 0.0000000 | 0.375 | 0.0000000 | 0.2000000 |  |  |  |
| 6 | 6 | 0 | 8 | 1 | 0.8571429 | 1.000 | 1.0000000 | 0.9333333 | 1 | 5 | 3 |
| 6 |  |  |  |  | 0.1428571 | 0.375 | 0.1666667 | 0.2666667 |  |  |  |
| 7 | 7 | 0 | 8 | 0 | 1.0000000 | 1.000 | 1.0000000 | 1.0000000 | 2 | 5 | 3 |
| 5 |  |  |  |  | 0.2857143 | 0.375 | 0.2857143 | 0.3333333 |  |  |  |
| 8 | 7 | 1 | 7 | 0 | 1.0000000 | 0.875 | 0.8750000 | 0.9333333 | 3 | 5 | 3 |
| 4 |  |  |  |  | 0.4285714 | 0.375 | 0.3750000 | 0.4000000 |  |  |  |
| 9 | 7 | 2 | 6 | 0 | 1.0000000 | 0.750 | 0.7777778 | 0.8666667 | 4 | 5 | 3 |
| 3 |  |  |  |  | 0.5714286 | 0.375 | 0.4444444 | 0.4666667 |  |  |  |
| 10 | 7 | 3 | 5 | 0 | 1.0000000 | 0.625 | 0.7000000 | 0.8000000 | 4 | 6 | 2 |
| 3 |  |  |  |  | 0.5714286 | 0.250 | 0.4000000 | 0.4000000 |  |  |  |
| 11 | 7 | 4 | 4 | 0 | 1.0000000 | 0.500 | 0.6363636 | 0.7333333 | 4 | 7 | 1 |
| 3 |  |  |  |  | 0.5714286 | 0.125 | 0.3636364 | 0.3333333 |  |  |  |
| 12 | 7 | 5 | 3 | 0 | 1.0000000 | 0.375 | 0.5833333 | 0.6666667 | 5 | 7 | 1 |
| 2 |  |  |  |  | 0.7142857 | 0.125 | 0.4166667 | 0.4000000 |  |  |  |
| 13 | 7 | 6 | 2 | 0 | 1.0000000 | 0.250 | 0.5384615 | 0.6000000 | 5 | 8 | 0 |
| 2 |  |  |  |  | 0.7142857 | 0.000 | 0.3846154 | 0.3333333 |  |  |  |
| 14 | 7 | 7 | 1 | 0 | 1.0000000 | 0.125 | 0.5000000 | 0.5333333 | 6 | 8 | 0 |
| 1 |  |  |  |  | 0.8571429 | 0.000 | 0.4285714 | 0.4000000 |  |  |  |
| 15 | 7 | 8 | 0 | 0 | 1.0000000 | 0.000 | 0.4666667 | 0.4666667 | 7 | 8 | 0 |
| 0 |  |  |  |  | 1.0000000 | 0.000 | 0.4666667 | 0.4666667 |  |  |  |

### Plot the results

graph_corr_annot () takes the output from get_confusionMatrix() to plot the final result, where precisions were plotted against the ranked pairs of mutants. Enrichment for sensitivity, specificity, precision, and accuracy could be compared with the dotted line, which is the negative control. For example:

> confusionMatrix_obj <-get_confusionMatrix_and_metrics (new,” same_annot”,” similarity”,seed=103)

> metric=“precision”; similarity=“pcc”; subset=dim(confusionMatrix_obj)[1]; cols=“blue”; ylim=c(0,1); lwd=1 # set graphing parameters

> graph_corr_annot(confusionMatrix_obj, metric, similarity, subset, cols, x_lab=““, ylim=c(0,1.05), lwd)

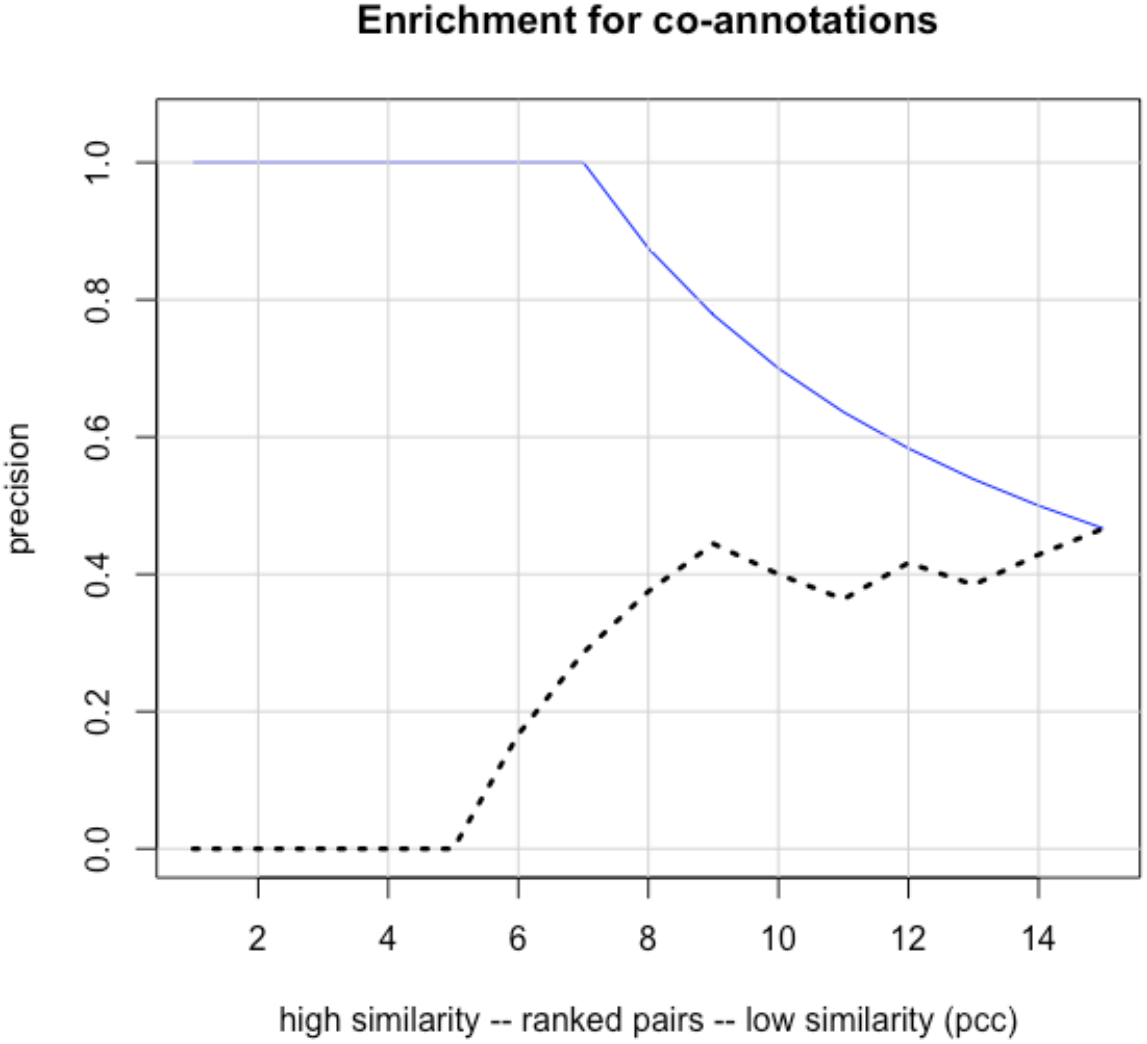

>metric=“sensitivity” # use another metric for the y axis

>graph_corr_annot(confusionMatrix_obj, metric, similarity, subset, cols, x_lab=““, ylim=c(0,1.05), lwd)

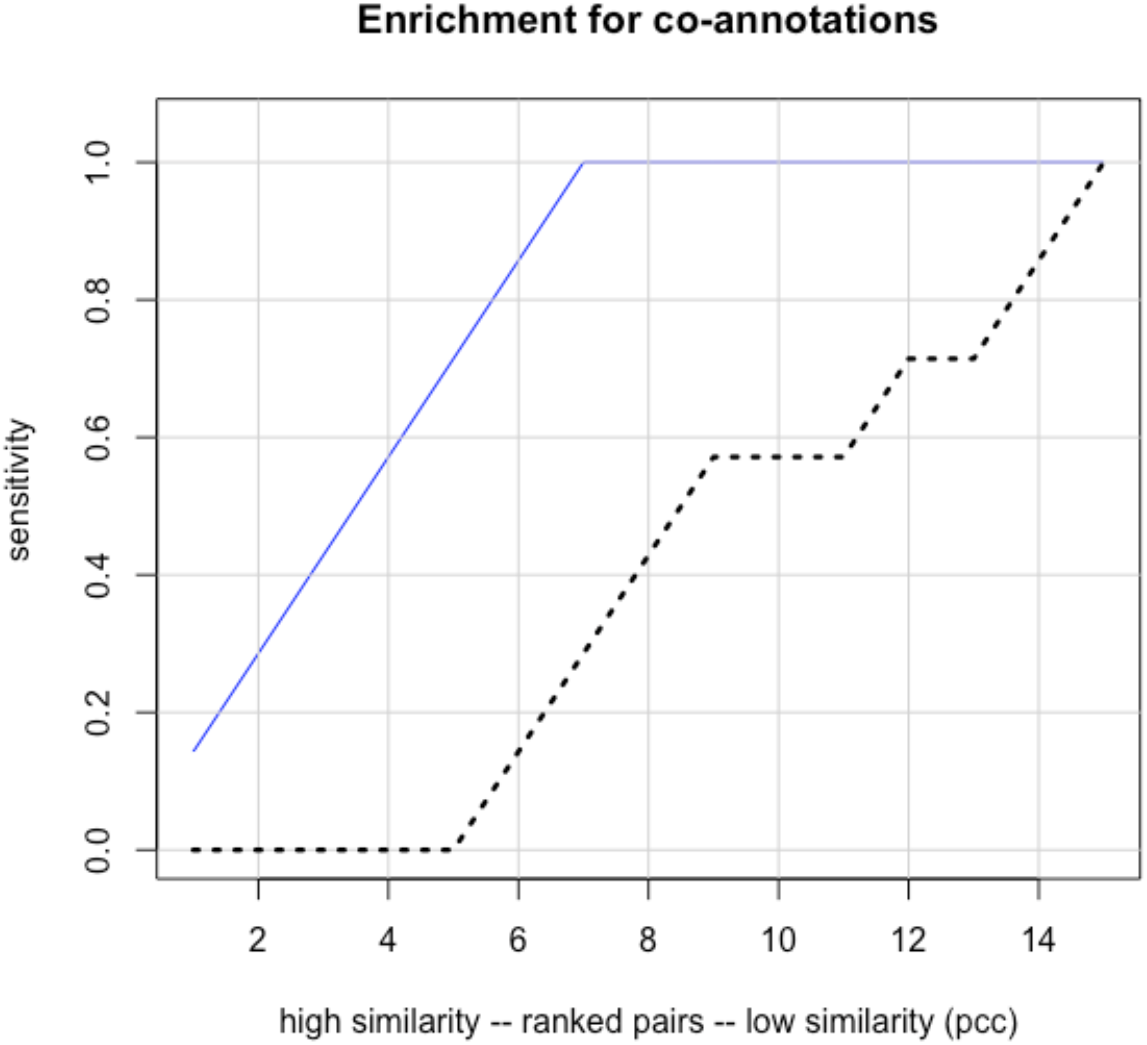

### Helper functions

The following 10 functions are intended to speed up the phenotype data curation process:

any_incomplete() checks if the input matrix, dataframe or datatable contains any NA, NAN, NULL or ““ (empty string). For example:

> incomplete_phenotype_data<-phenotype_data

> incomplete_phenotype_data[1:2, 1:2]<-NA #introduce some NA

> incomplete_phenotype_data

|  | STREPTOMYCIN.0.05 | SUCCINATE | SULFAMONOMETHOXINE.100 | TAUROCHOLATE.0.1 | TAUROCHOLATE.0.5 |
| --- | --- | --- | --- | --- | --- |
| ECK3997-purD | NA | NA | 3.33 | -13.46 | - |
| 13.79 |  |  |  |  |  |
| ECK0516-purE | NA | NA | 2.63 | -8.27 | - |
| 9.81 |  |  |  |  |  |
| ECK3763-ilvD | -1.41 | -11.72 | 0.34 | -0.69 | 0.32 |
| ECK3766-ilvC | -0.58 | -8.53 | -0.81 | 0.06 | 0.51 |
| ECK3762-ilvE | -0.01 | -11.93 | 0.02 | -0.68 | - |
| 0.46 |  |  |  |  |  |
| ECK3764-ilvA | -0.15 | -11.55 | -0.84 | -0.04 | - |
| 0.19 |  |  |  |  |  |

> any_incomplete(incomplete_phenotype_data)

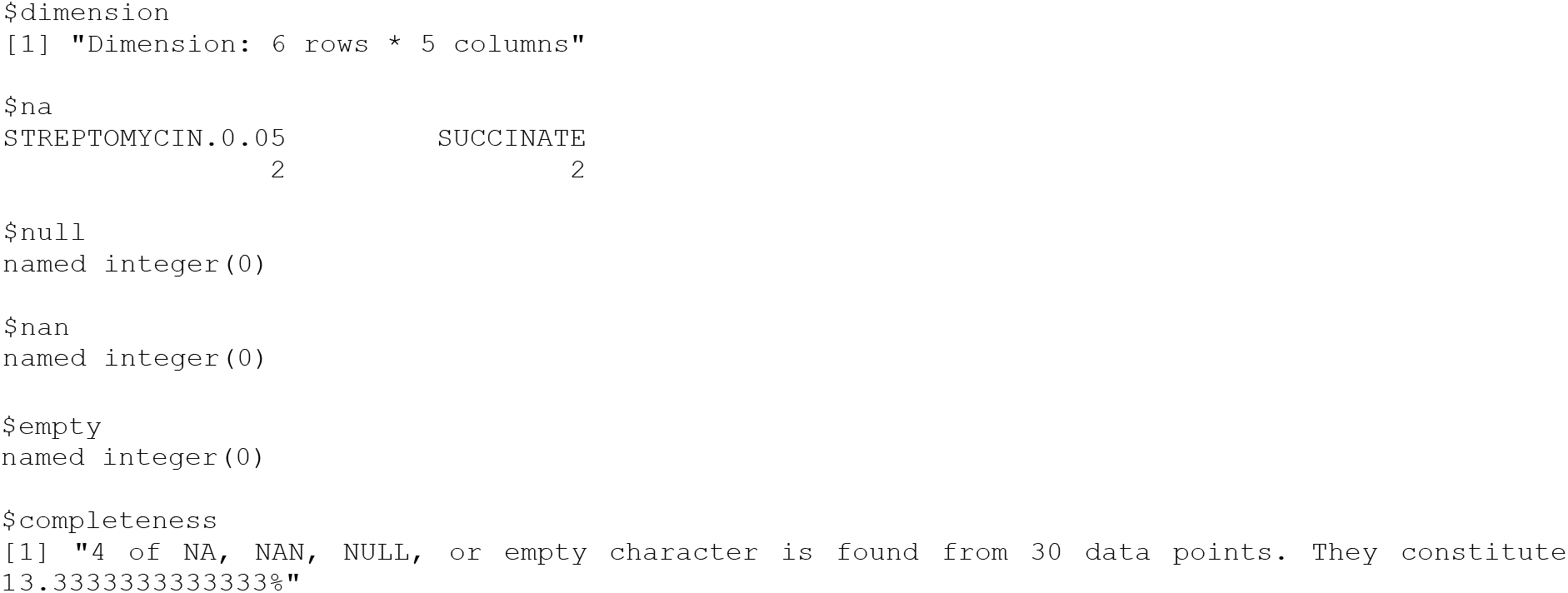

filter_table() filters the input matrix, dataframe or datatable so that all rows and columns with NA/NAN/NULL/” “are removed. For example:

> filter_table(incomplete_phenotype_data)

|  | SULFAMONOMETHOXINE.100 | TAUROCHOLATE.0.1 | TAUROCHOLATE.0.5 |
| --- | --- | --- | --- |
| ECK3997-purD | 3.33 | -13.46 | -13.79 |
| ECK0516-purE | 2.63 | -8.27 | -9.81 |
| ECK3763-ilvD | 0.34 | -0.69 | 0.32 |
| ECK3766-ilvC | -0.81 | 0.06 | 0.51 |
| ECK3762-ilvE | 0.02 | -0.68 | -0.46 |
| ECK3764-ilvA | -0.84 | -0.04 | -0.19 |

graph_table() takes a matrix, dataframe or a table as the input and represents it using a heatmap. It also deals with continuous/categorical/mixed variables. For example:

> graph_table(phenotype_data)

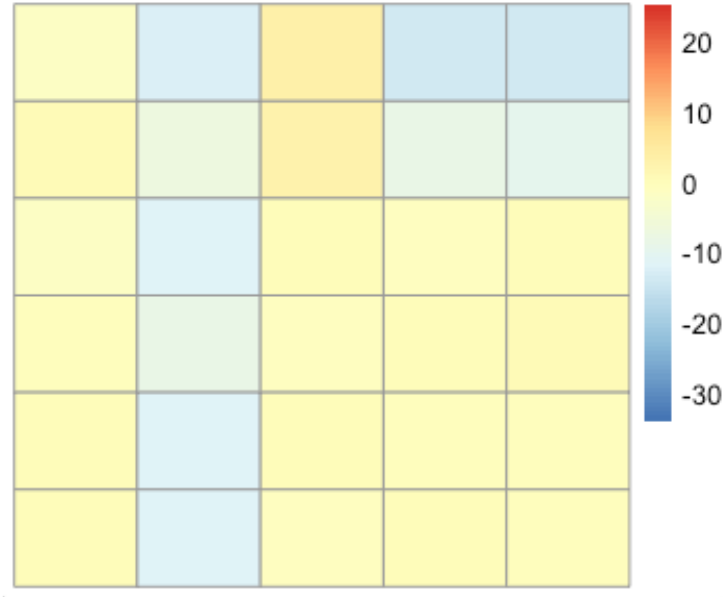

> graph_table(ter_phenotype)

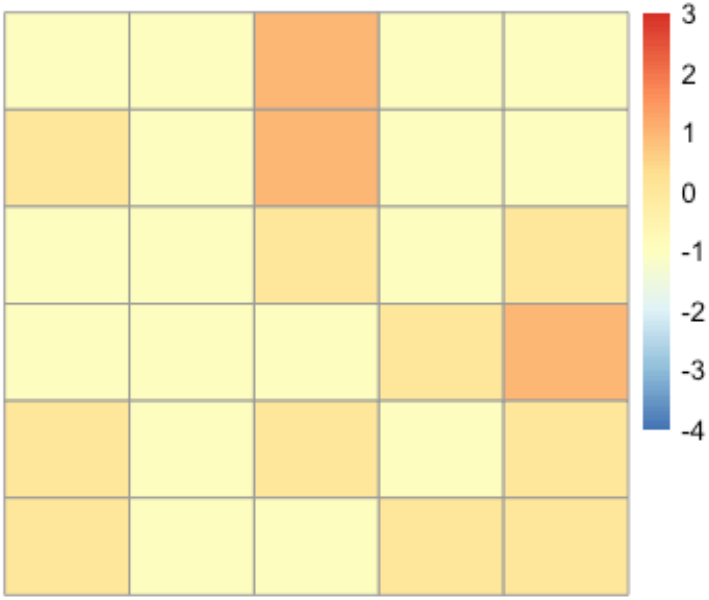

checkDuplicates_vect() checks if an input vector has duplicates. If so, it will return the frequency table. Otherwise, the text “Everything in this vector is unique” will be returned. For example:

> my_vector <-c(1,1,2,2,3)

> checkDuplicates_vect(my_vector)

[1] “Some duplicates are found:”

vect

1 2 3

2 2 1

> my_vector <-c(1,2,3)

> checkDuplicates_vect(my_vector)

[1] “Everything in this vector is unique”

change_names () changes row names or column names of a matrix, dataframe or datatable based on another matrix/dataframe/datatable. For example:

> new_col_names <-c(“0.05 μg/ml streptomycin”,

“0.3% succinate”,

“100 μg/ml sulfamonomethoxine”,”0.1% taurocholate”,”0.5% taurocholate “)

> change_names(rowOrCol=“col”, phenotype_data, matrix(c(colnames(phenotype_data), new_col_names), ncol=2, byrow=FALSE))

|  | 0.05 µg/ml streptomycin | 0.3% succinate | 100 µg/ml sulfamonomethoxine | 0.1% taurocholate | 0.5% taurocholate |
| --- | --- | --- | --- | --- | --- |
| ECK3997-purD | -1.48 | -12.30 | 3.33 | -13.46 | -13.79 |
| ECK0516-purE | 0.43 | -6.75 | 2.63 | -8.27 | -9.81 |
| ECK3763-ilvD | -1.41 | -11.72 | 0.34 | -0.69 | 0.32 |
| ECK3766-ilvC | -0.58 | -8.53 | -0.81 | 0.06 | 0.51 |
| ECK3762-ilvE | -0.01 | -11.93 | 0.02 | -0.68 | -0.46 |
| ECK3764-ilvA | -0.15 | -11.55 | -0.84 | -0.04 | -0.19 |

melt_similarity() takes a similarity matrix or a distance object as the input and converts it to a long form dataframe, with an option to sort the molten dataframe by the 3^rd^ column (the numeric column). For example:

> dist(phenotype_data) # the distance object of interest

|  | ECK3997-purD | ECK0516-purE | ECK3763-ilvD | ECK3766-ilvC | ECK3762-ilvE |
| --- | --- | --- | --- | --- | --- |
| ECK0516-purE | 8.815730 |  |  |  |  |
| ECK3763-ilvD | 19.272945 | 13.906973 |  |  |  |
| ECK3766-ilvC | 20.480305 | 13.853281 | 3.575766 |  |  |
| ECK3762-ilvE | 18.822093 | 13.374180 | 1.647726 | 3.749973 |  |
| ECK3764-ilvA | 19.615726 | 13.989089 | 1.921328 | 3.131485 | 1.177327 |

> melt_similarity(dist(phenotype_data))

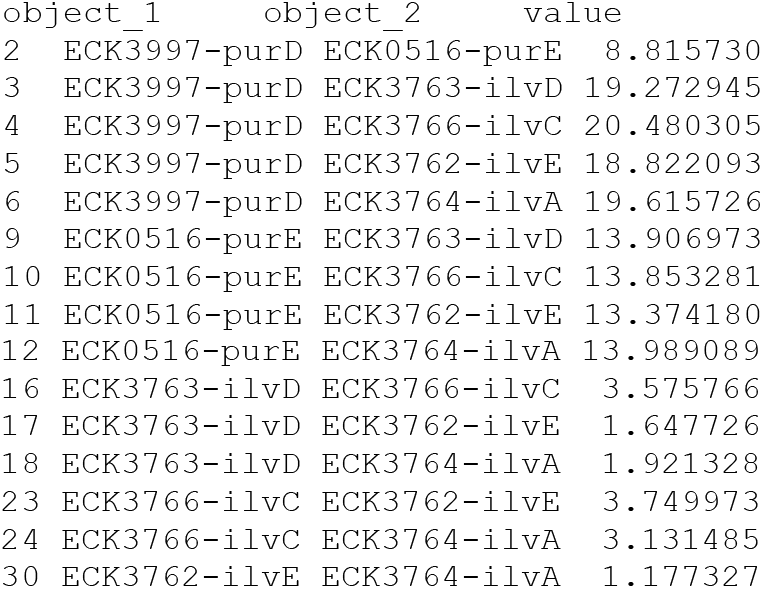

When compared with reshape2::melt(), the differences are: 1. melt_similarity() Can take a distance object as the main input 2. When a matrix is used as the main input, it has to be a similarity matrix 3. melt_similarity() remove the diagonal elements and the duplicated pairs that share the same similarity.

convert_table() takes a matrix, dataframe or a datatable as the input and converts the types of elements to a designated type. For example:

> str(phenotype_data) remove_NA()

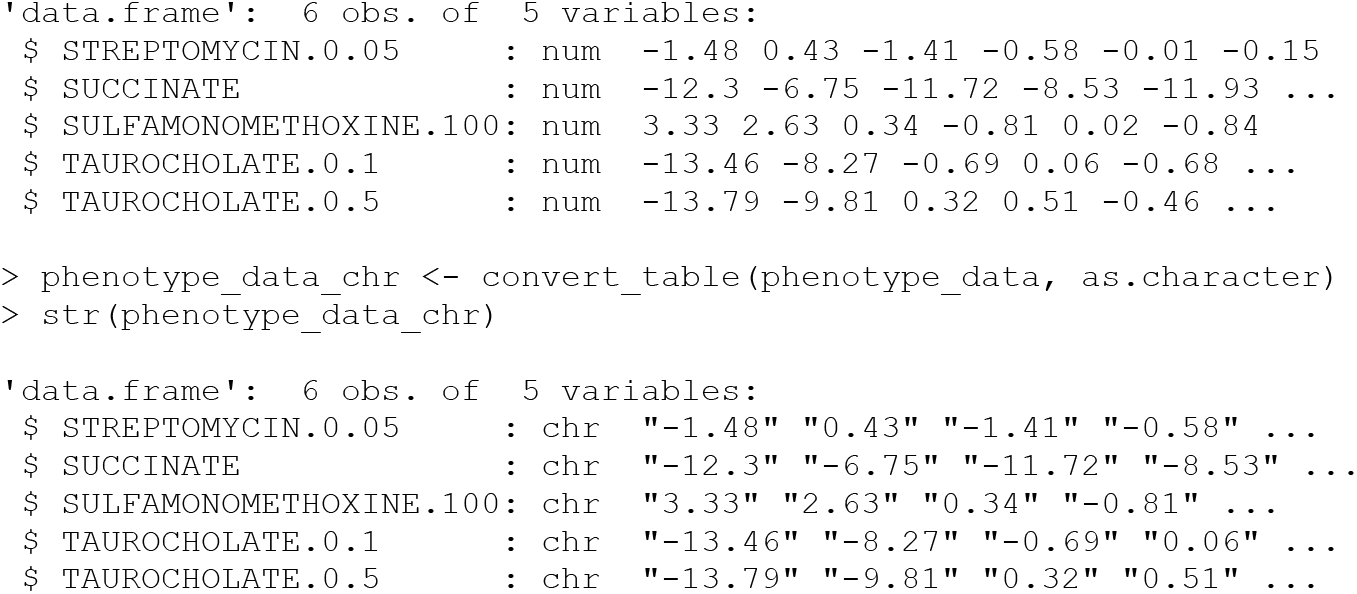

removes NA from an R vector object. For example:

> remove_NA (c(1,2,3,4,NA,NA))

[1] 1 2 3 4

### Summary

The microbialPhenotypes package has implemented a pipeline to systematically parse and analyze high-throughput phenotype screens, with many functions as well as an example using a published *E. coli* dataset (Nichols et al., 2011). Although the motivation for this software is to process microbial phenotype data, we expect its usability to be easily extended to multivariate dataset with distinct annotation sets.

## Acknowledgements

This work was supported by a grants from the National Science Foundation Division of Biological Infrastructure [1458400] and the National Institutes of Health [R01GM089636, U41HG008735]. PW wrote the package and drafted the manuscript. DS, JH supervised the project as well as testing the usability of the code.

